# Mathematical modeling of the role of α-synuclein and dopamine in cell death in Parkinson’s disease provides the molecular basis to the toxicity of different point mutations

**DOI:** 10.1101/714600

**Authors:** Gourvendu Saxena, Utkarsh Khandelwal, Mukesh Doble

**Affiliations:** NUS Yong Loo Lin School of Medicine, Singapore-119228; Department of Biotechnology, Indian Institute of Technology Madras, Chennai, India-600036

**Keywords:** NAC peptide, Neurodegeneration, Oxidative stress, Parkinson’s disease, Reactive oxygen species, α-synuclein

## Abstract

Different types of α-synuclein and non-β-amyloid component (NAC) peptides have been shown to induce cell death, with varying degree of toxicity, in various in vitro experiments. Oxidative stress has also been associated and proved to be involved in the pathogenesis of neuronal cell death in Parkinson’s disease. Oxidative stress has been shown to accelerate the aggregation of α-synuclein in vitro and in resent studies α-synuclein has been shown to increase oxidative stress. Thus it seems like a vicious cycle, one promoting the other.

In this present work we have modeled the α-synuclein pathway to increase cytoplasmic Dopamine concentration, and thereby increasing the Reactive Oxygen Species (ROS) level of the cell, which consequently results in cell death. This model relates the α-synuclein concentration with the fractional cell survival and provides insight of crucial reaction(s) of α-synuclein which promote cell death. It predicts the toxicity of the type of α-synuclein and also explains the pattern of cell death with increasing concentration of α-synuclein. First we modeled a part of the pathway i.e. from dopamine to cell death. The results were compared with experimental data available for PC12 neuronal cell line. Then modeling of full pathway was done and the results were compared with experimental data available for Human neuroblastoma SH-SY5Y cells. It is predicted from this model that higher the auto catalysis of dopamine, higher is the cell death. Interestingly, the model predicts that NAC (1-18) not only hinders the vesicles coming from Endoplasmic Reticulum, to fuse with Golgi bodies, but also reduces the synthesis of Dopamine and the formation of vesicles from Endoplasmic Reticulum. The model is generalized and can predict the toxicity of any protein which impedes the early secretary pathway in dopaminergic cells and also the cell survival pattern with increasing concentration of the protein.

## INTRODUCTION

Parkinson’s disease (PD) is the second most common neurodegenerative disorder (Forman et al., 2004) and affects nearly 1% of the population who are above sixty years old. The disease is caused due to extensive loss of Dopaminergic neurons in substantia nigra pars compacta in basal ganglia of the brain. Evidences from the past research have shown the central role of protein called α-synuclein in the pathogenesis. The hallmark of PD is the formation of Lewy bodies and cellular inclusions (Spillantini et al., 1998). The missense mutations (A53T, A30P, E46K) are also associated with the pathogenesis of PD. Duplication or triplication in several families are also linked to early onset of familial PD (Singleton et al., 2003).

α -synuclein is a highly conserved protein with 140 amino acids residues and it is abundant in presynaptic terminals (Iwai et al., 1995). Originally it was identified as the precursor of the non-β-amyloid component (NAC) (Ueda et al., 1993). NAC is a 35 amino acids peptide comprising of amino acid 61-95 of the α-synuclein and has been identified as the second major constituent in the plaques of Alzheimer’s disease (AD) (Ueda et al., 1993). Mutation in the α-synuclein causes alteration in its amino acid sequence (A30P or A53T) in regions which are predicted to influence the secondary structure. The substitution may disrupt the structure of α-synuclein, rendering the protein more prone to self aggregation and hence disposition to Lewy bodies (Kruger et al., 1998). In a recent study it has been suggested that abnormal α-synuclein accumulation is likely to impede the early secretary pathway in many cell types (Susan et al., 2006).

Dopamine is synthesized in the cytoplasm and is pumped rapidly into the vesicles. Defects in the early secretary pathway may cause shortage of vesicles to sequester the Dopamine that is being synthesized in the cytoplasm. Unsequestered dopamine is unstable and oxidizes to generate Reactive Oxygen Species (ROS) producing hydrogen peroxide (Asanuma et al. 2003). More specifically, the inhibition of vesicular trafficking by α-synuclein may affect dopamine-producing neurons because neurotransmitters produced by the other neurons are less toxic than Dopamine (Susan et al., 2006). Post mortem studies provide ample evidence for increased oxidative stress and oxidative damage to lipids proteins and DNA in the nigra of PD patients (Alam et al., 1997). A leading hypothesis about the biochemical pathogenesis of PD centers around the formation of ROS leading to oxidative damage to nigral dopaminergic neurons (Jenner et al., 1998).

Despite extensive study little is known about the normal functioning of α-synuclein or how it contributes to the disease. In this present work, using mathematical modeling approach we have tried to elucidate one of the pathological pathway by which α-synuclein could cause cell death. We have also tried to find out the effect of generation of dopamine by the cell on its vulnerability towards α-synuclein toxicity. The model predictions are compared with the experimental observations.

## METHODS AND PROCEDURE

### Model Description

In numerous experiments α-synuclein and NAC peptides have been demonstrated to self aggregate in proto fibrils and amyloid like fibrillar and β-sheet conformations (Conway et al., 2000). The proto fibril is an intermediate species between the α-synuclein monomer and the elongated fibrils. Based upon observation of the disappearance of the monomer, it could be assumed that both the A53T and the A30P mutants appear to form proto fibrils faster than WT α-synuclein, although the A53T variant shows more accelerated fibril formation. Metals like iron and Cu (II) have been reported to induce α-synuclein aggregation, in the presence of H_2_O_2_ (Conway et al., 2000). The intermediate proto fibril is the pathological species which causes PD. The aggregation of α-synuclein and the effect of different reactive oxygen species (ROS) have already been modeled (Raichur et al., 2006). Here we have taken pathological α-synuclein as our starting point for modeling, which leads to cell death, thereby relating α-synuclein concentration with number of dead cells (Fig 1). The pathway shown in figure 1 is created based on information provided in the literature and it is in consistence with previous studies (Susan et al., 2006, Takafumi Hasegawa et al., 2006).

**Figure 1:**
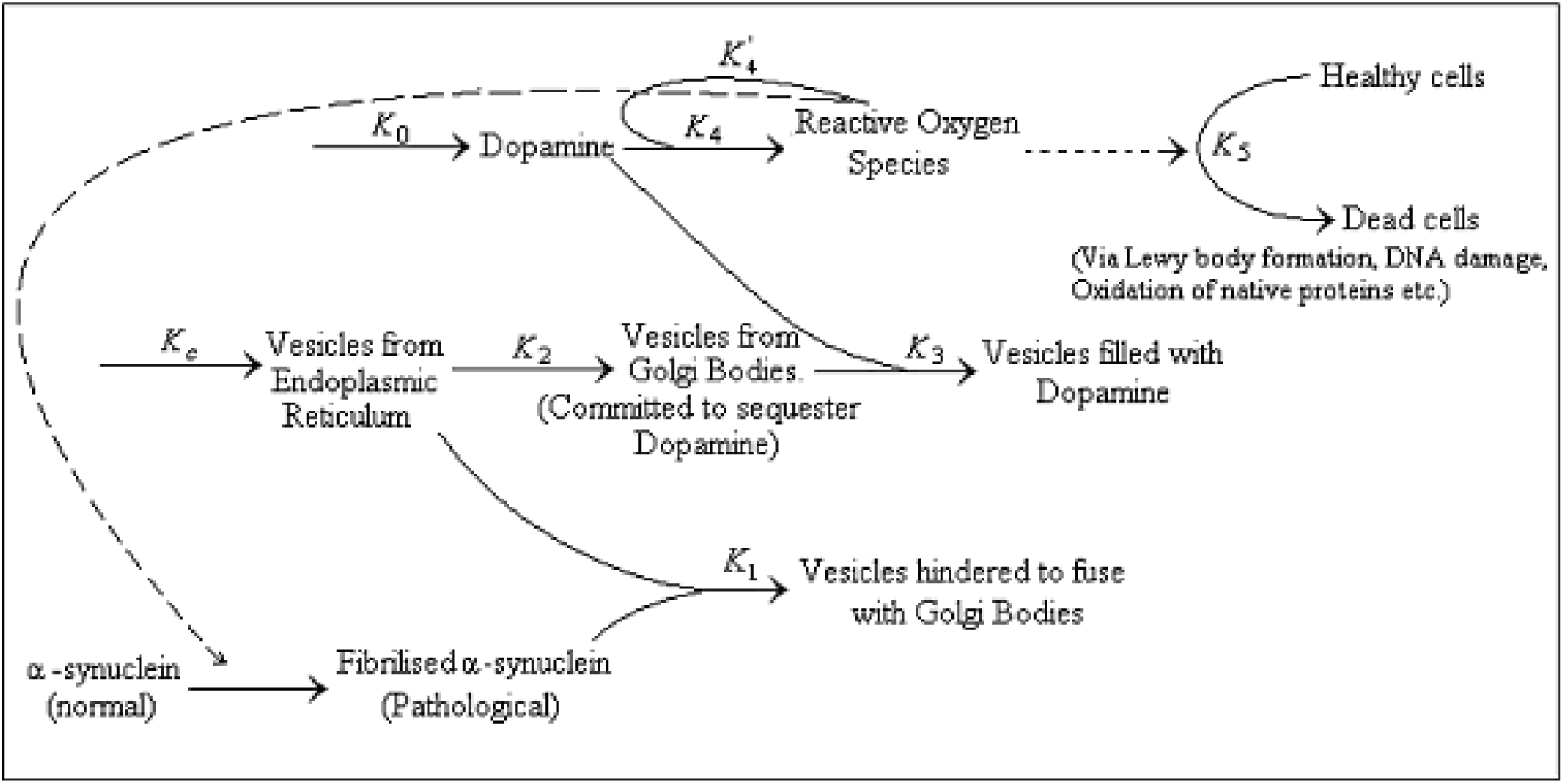
Map of α-synuclein pathway to cell death. Pathological α-synuclein hinders the early secretary pathway, due to which there is a shortage of vesicles to sequester dopamine formed in the cytoplasm. This results in the high intracytoplasmic concentration of dopamine. Dopamine is oxidized to form ROS causing cell death.

The vesicles from Endoplasmic Reticulum (ER) have been assumed to emerge with zero order kinetics because the surface area of ER does not reduce even after emergence of vesicles. These vesicles fuse to Golgi Bodies (GB) and then are committed to sequester dopamine formed in the cytoplasm. The freely formed dopamine in the cytoplasm is pumped into the vesicles coming from GB, thus sequestering dopamine in vesicles. The α-synuclein hinders the early secretary pathway of the cell, i.e. fusion of vesicles emerging from ER to GB. This α-synuclein mediated defect in the early secreted pathway decreases the steady state concentration of vesicles committed to sequester dopamine, thus there is shortage of volume to store free dopamine in cytoplasm (Susan et al., 2006). So the free dopamine starts building up in the cytoplasm, which starts oxidizing to form the Reactive Oxygen Species (ROS) in the form of H_2_O_2_. Once ROS starts accumulating it further accelerates the oxidation of dopamine to ROS in an autocatalytic manner thereby increasing the concentration further. (Yulia Y. Tyurina, 2006). Hence, dopaminergic cells are more prone to α-synuclein and dopamine toxicity. This ROS accumulation ultimately causes cell death through various means which includes DNA damage, lipid and protein peroxidation, cytoskeleton disorganization etc (Blum D. et al., 2001).

This cell death pathway can be modeled as a time dependent phenomenon as shown below.

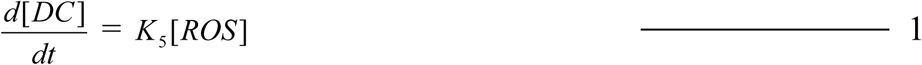

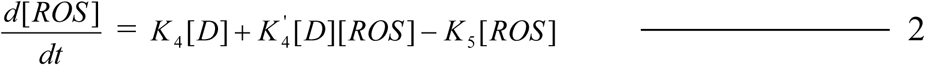

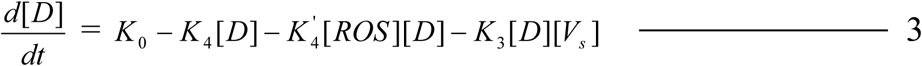

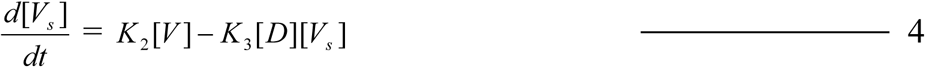

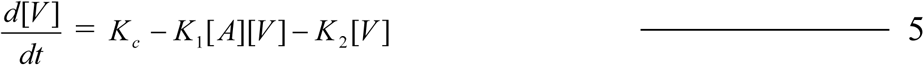

Where:

[A] = Fibrilised α**-**Synuclein Concentration
[DC] = Concentration of Dead Cells.
[D] = Concentration of Dopamine.
[D_0_] = Initial Dopamine Concentration.
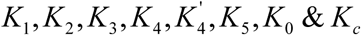 are rate constants for different reactions (Figure.1)
[ROS] = Reactive Oxygen Species Concentration
t_f_ = Time of induction.
[V] = Concentration of Vesicles emerging from endoplasmic Reticulum
[V_s_] = Concentration of Vesicles emerging from Golgi body.

If reactions 2, 3, 4 & 5 are assumed to be reaching steady state the relationships between the cell death as a function of dopamine and as a function of fibrilised α-synuclein respectively are obtained as follows:

(Appendix I gives a detailed derivation for equations 6 & 7)

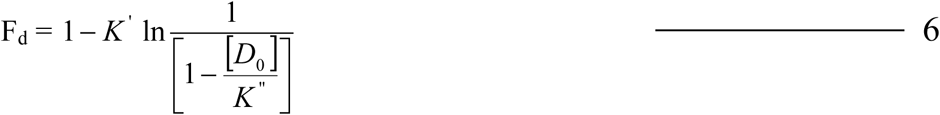

Equation 6 relates the fractional cell survival (F_d_) as a function of initial dopamine concentration ([D_0_]). This equation has two constants *K*′ & *K*″

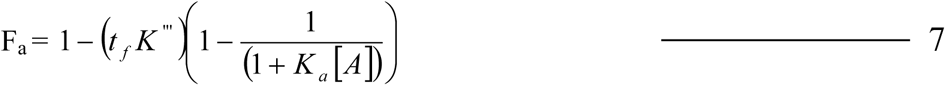

Equation 7 relates the fractional cell survival (F_a_) as a function of fibrilar α-synuclein concentration ([A]) after time, t_f_. This equation has two constants *K*‴ & *K*_*a*_.

The justification for constructing such a model is that, it can account for toxicity of different types of pathological α-synuclein and also explain the characteristics of fractional cell survival as a function of α-synuclein concentration similar to a dose response relationship. The studies pertaining to over expression of various kinds of α-synuclein, NAC peptide etc giving rise to cell death cannot be predicted by this model because the model requires fractional cell survival as a function of α-synuclein concentration, which remains unavailable in experiments relating to over expression of α-synuclein.

α-synuclein has been shown to induce neuronal cell death by Rab5A dependent endocytosis (Jee Young Sung et al., 2001). Findings, that α-synuclein is secreted into Cerebrospinal fluid (Borghi et al., 2000) and freshly prepared or aggregates of α-synuclein induce cell death (El-Agnaf et al., 1998), underline the importance of such a model. Further, it has been shown in the past that over expression of tyrosinase induces apoptosis and co-expression of wild type or A53T mutant human α-synuclein with tyrosinase further exacerbates cell death in SH-SY5Y cells (Takafumi Hasegawa at el., 2006). The proto fibrilised α-synuclein, inside the cell has been proved to interfere with ER-GB trafficking, i.e. it manages to drag away Rab1, which helps in the fusion of vesicles coming from ER to GB (Susan et al., 2006). Hence it is suggested that there may be a shortage of vesicles and Vesicular Monoamine Transporter 2 (VMAT 2) required to pump free dopamine into the vesicles. Due to the shortage of vesicles, the free dopamine concentration starts building up in the cytoplasm which generates the ROS. ROS further accelerates α-synuclein aggregation and dopamine oxidation, the latter in turn generates more ROS. This elevated ROS causes cell death. Further, α-synuclein-dopamine adducts stabilizes protofibrils and inhibits fibrilization, thus stabilizing the α-synuclein oligomers in the proto fibriler state (Conway et al. 2001). There are reports which give evidence of increased levels of ROS in the cells when α-synuclein is over expressed or mutant versions of α-synuclein are expressed. (Eunsung Junn and Maral Mouradian, 2002) In this current work, the net effects of pathological states are taken into consideration.

For example, in the normal healthy cells it is assumed that there will be no accumulation of free dopamine in the cytoplasm. Thus, there is no extra free radical scavenging machinery available for extra ROS that is generated in the diseased cells. In this model the ROS scavenging machinery has not been included. Also α-synuclein is provided in excess to the cells from out side, thus there is no (or negligible) control of cell’s machinery on its cytoplasmic α-syunuclein concentration. Cell’s Ubiquitin proteosome complex mediated α-synuclein degradation has also been assumed to have negligible effect on cytoplasmic α-synuclein concentration for the same reason.

The constants in the model 6 and 7 can be determined by fitting reported experimental data. Addition of radical scavenger or antioxidant such as glutathione is found to decrease the fractional cell death (Blum D. et al., 2001), probably by quenching the ROS. This can be simulated using one more reaction step as shown in figure 2. This leads to an equation for fractional cell survival as

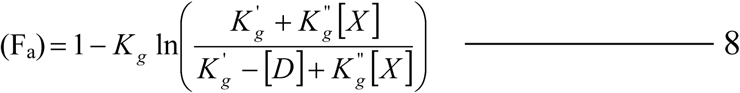

Where

[*X*] = Concentration of anti-oxidant.
[*D*] = Concentration of Dopamine.
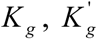 and 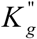 are the constants.

**Figure 2:**
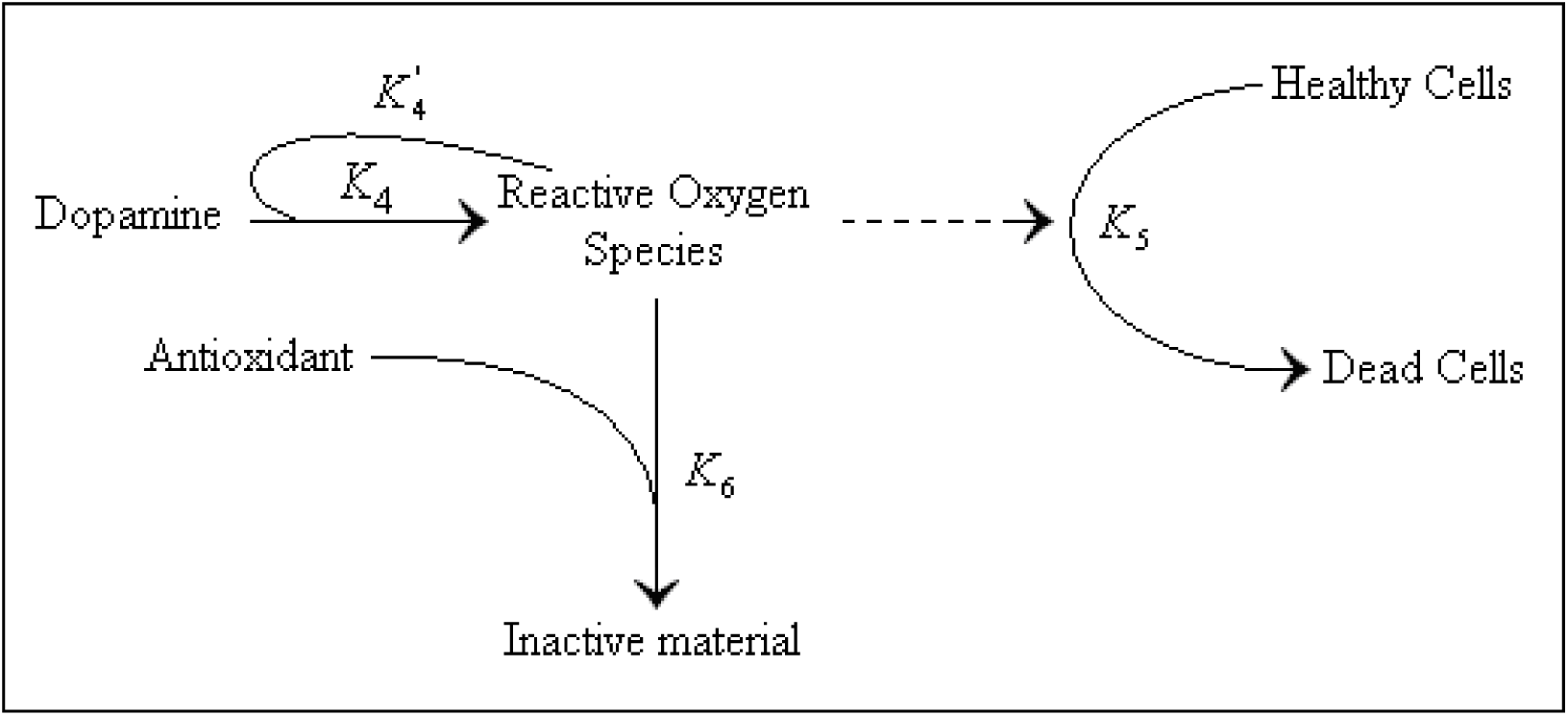
Effect of antioxidant on ROS concentration and cell death

## RESULTS

Figures 3 and 4 simulate the fractional cell survived as a function of dopamine concentration for PC12 neuronal cell line. The figures compare the model prediction with experimental data reported in the literature (Tyurina et al., 2006). The estimated *K*′ for the two data sets are 0.414 and 0.283 respectively whereas *K*″ is found to be 58 and 85 µ mol respectively. The constant 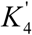 is an indication of the extent of autocatalysis. Higher the value of 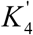, higher is the extent of autocatalysis and higher is the generation of ROS. This means that lower the value of 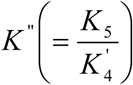 higher will be the cell death. Figure 5 shows the effect of decreasing *K*″on the fraction of cell survival for varying dopamine concentration.

**Figure 3:**
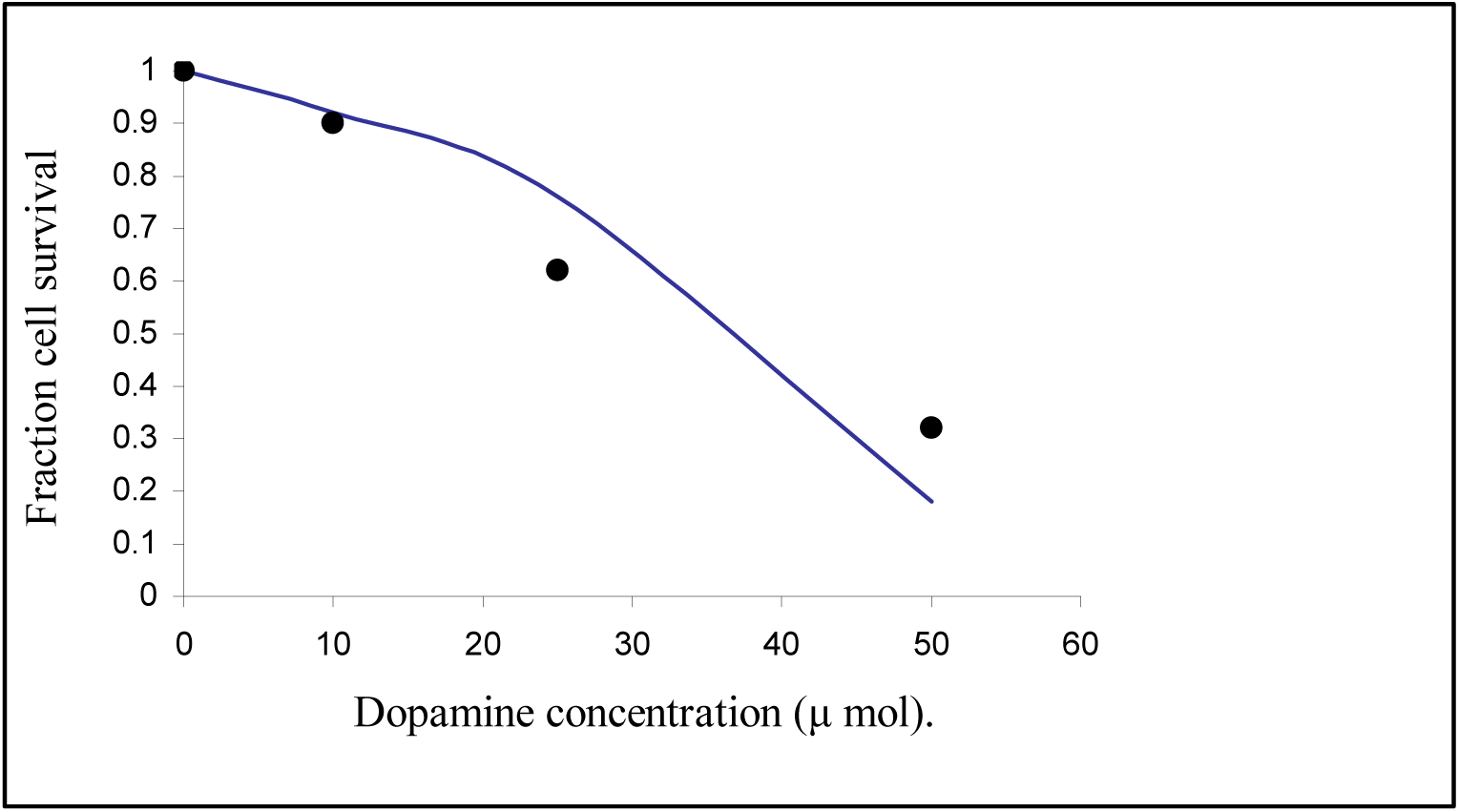
Comparison of results of dopamine concentration Vs cell death model with the experimental data (Tyurina et al., 2006). Cell line used is the neuronal PC12/mock. (Points = Data; Continues line = Model)

**Figure 4:**
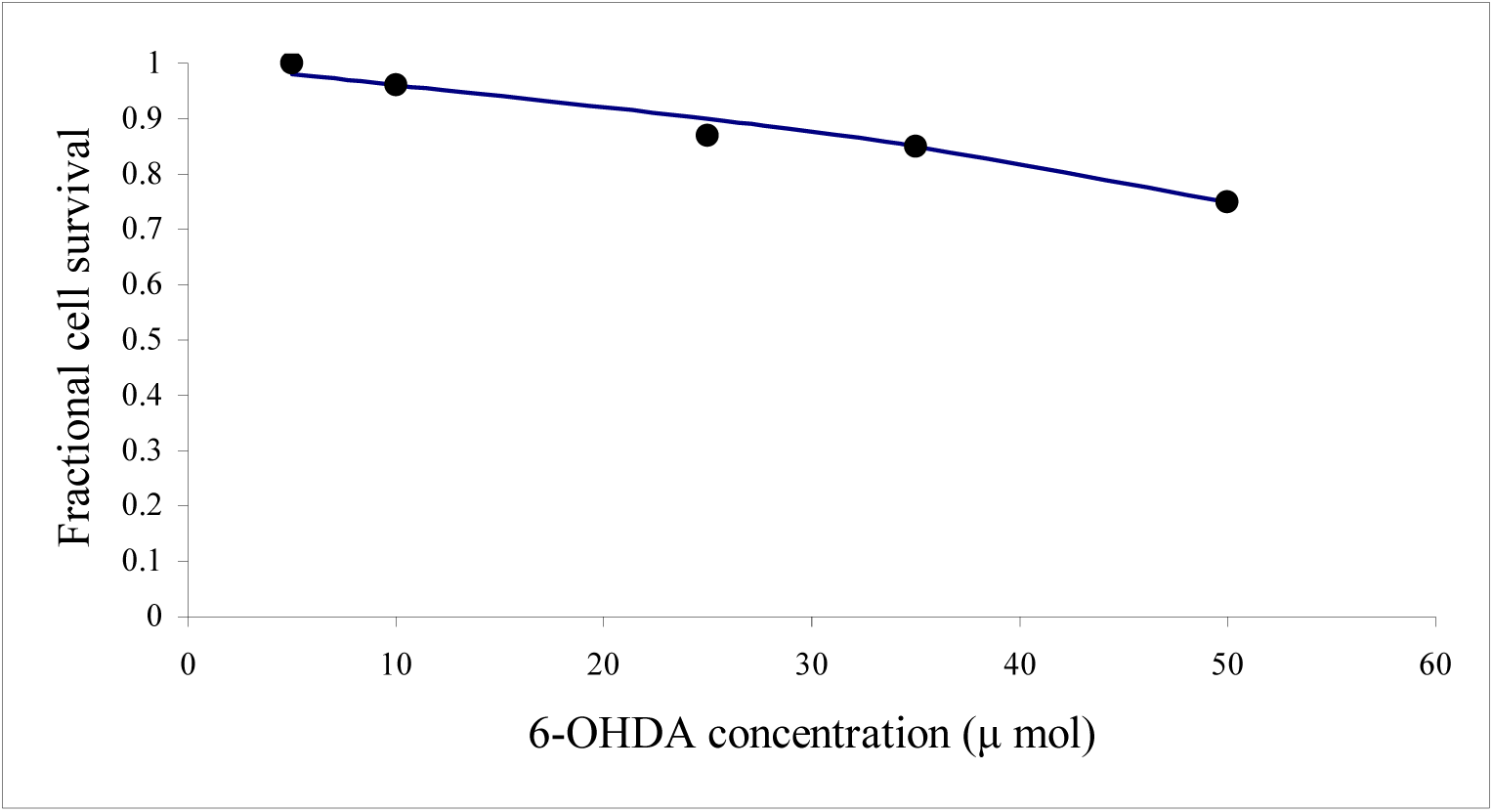
Comparison of results of 6-OHDA concentration Vs cell death model with the experimental data (Walkinshaw G. and Waters C. M., 1994). Cell line used is the neuronal PC12. (Points = Data; Continues line = Model)

**Figure 5:**
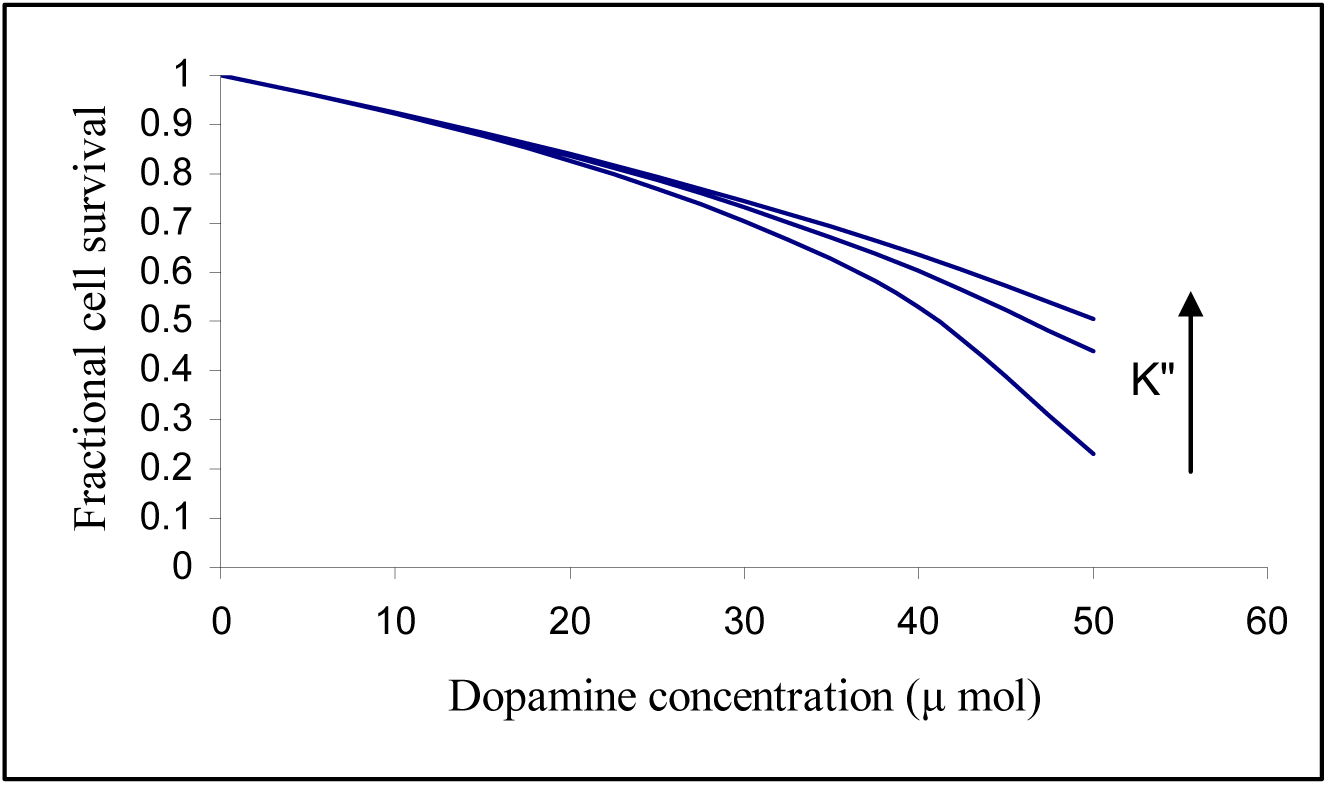
Simulation of dopamine Vs cell death model with different values of *K*″

Figure 6 shows the effect of adding an antioxidant such as Glutathione which scavenges the ROS on the cell death. The amount of ROS generated decreases due to the addition of antioxidant. The experimental data is taken from (Blum D. et al., 2001). A 10,000 µ mol of glutathione can prevent the cell death completely. The estimated 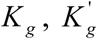 and 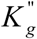 are 0.428, 150 and 0.05 respectively.

**Figure 6:**
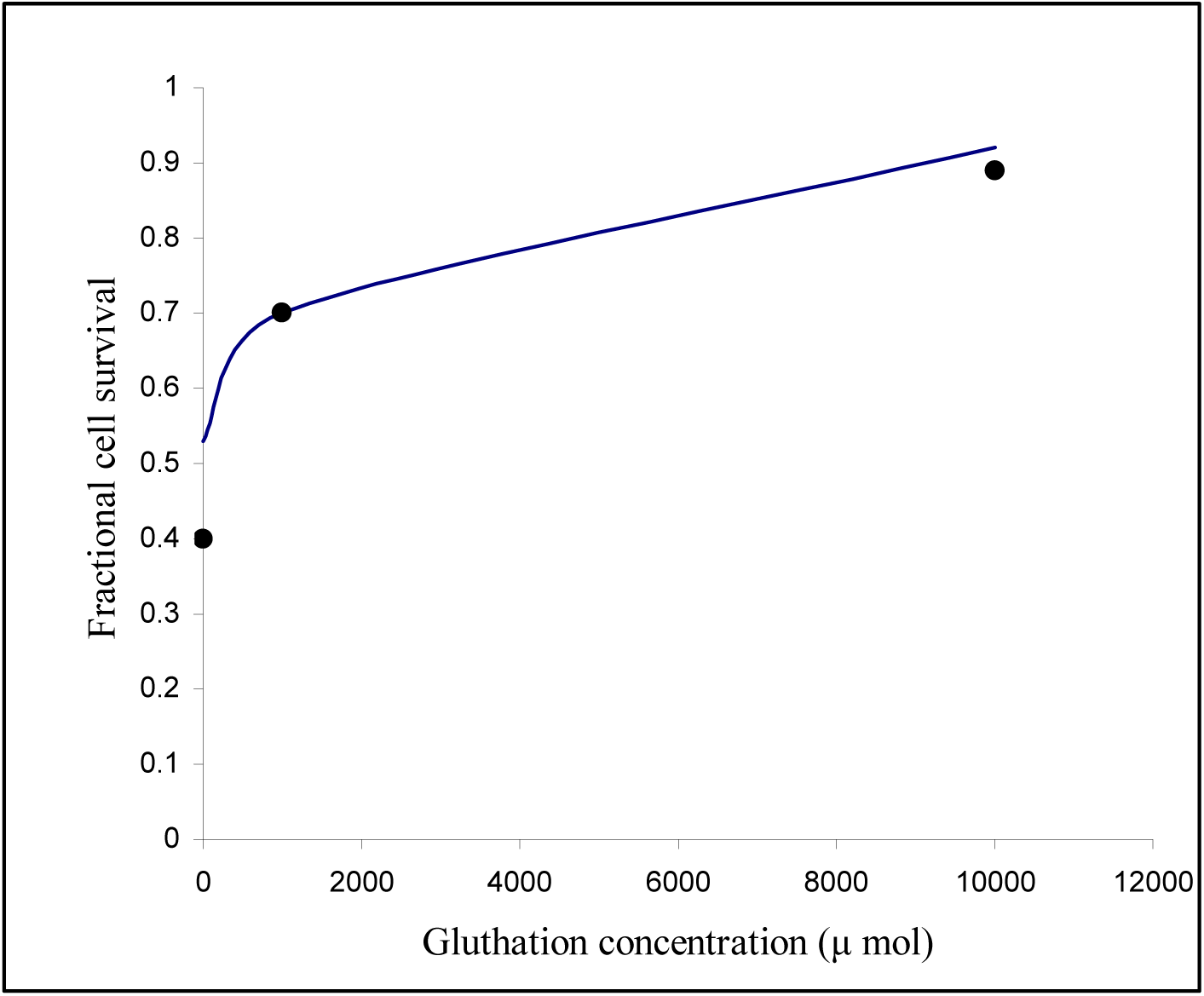
The effect of antioxidant (glutathione) on cell death when cells are kept at dopamine concentration of 100 μ mol. The experimental data (David Blum et al., 2001) is compared with model results. (Points = Data; Continues line = Model)

Figure 7 to 12 simulate the effect of increased level of aged α-synuclein on fraction of cell survival and compares with the experimental reported data. In all the cases the model is able to predict the observed behavior well. The type of α-synuclein considered in each of the experimental data is listed in table 1. 100% cell survive for small amounts of aged α-synuclein and, it suddenly drops when it reaches a threshold value, which happens to be 10^−1^ and10^−3^ μM for human dopaminergic neuroblastoma SH-SY5Y cells, exposed to aged α-synuclein (A30P) and to fresh NAC (1-18) peptide, respectively. The estimated values of *K*‴ and *K*_*a*_ are listed in table 1.

**Figure 7:**
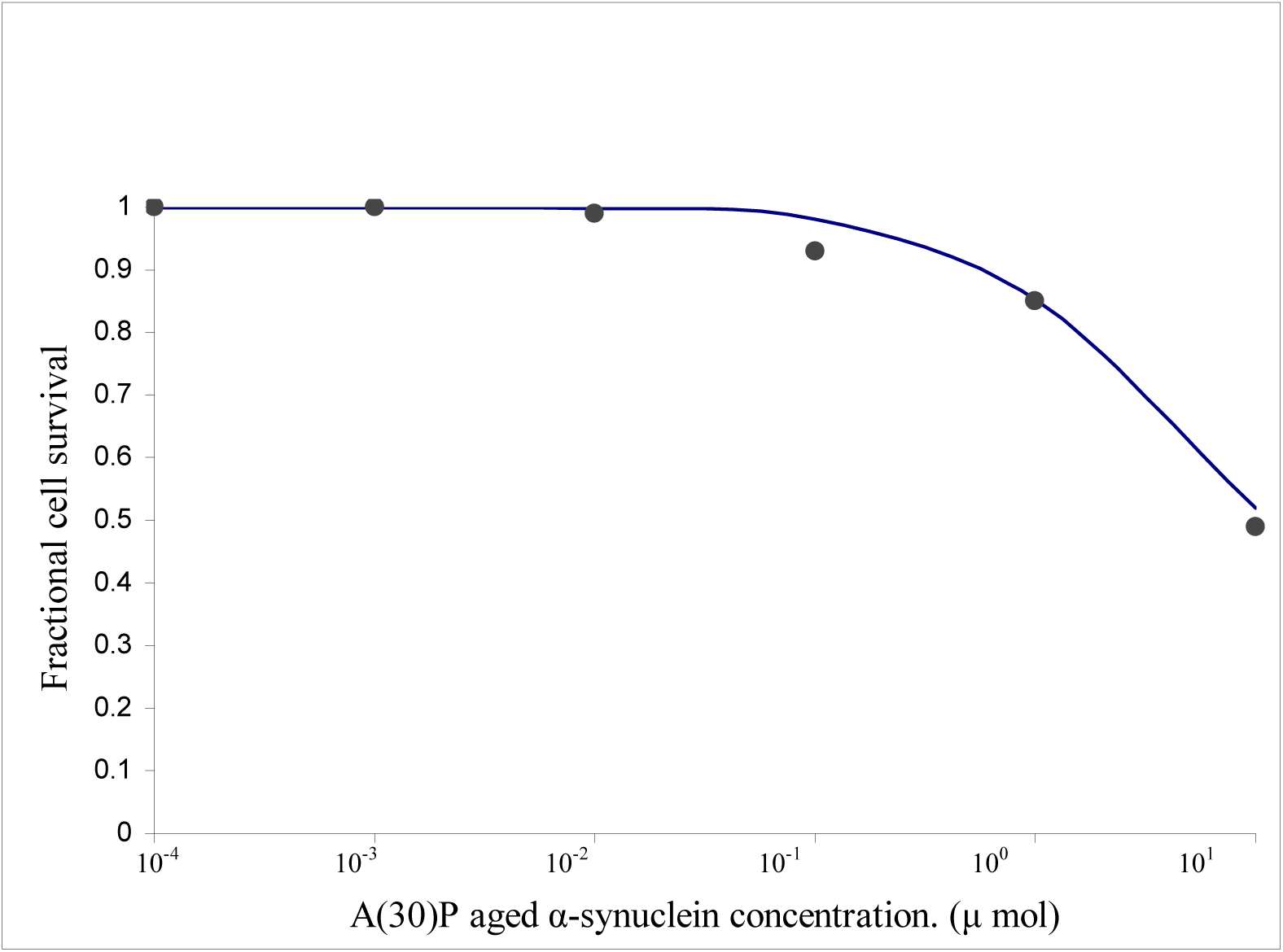
Human dopaminergic neuroblastoma SH-SY5Y cells were challenged with different concentrations of exogenous A (30) P aged α-synuclein. The experimental data (El-Agnaf et al., 1998) is compared with the results of α-synuclein Vs cell death model. (Points = Data; Continues line = Model)

**Figure 8:**
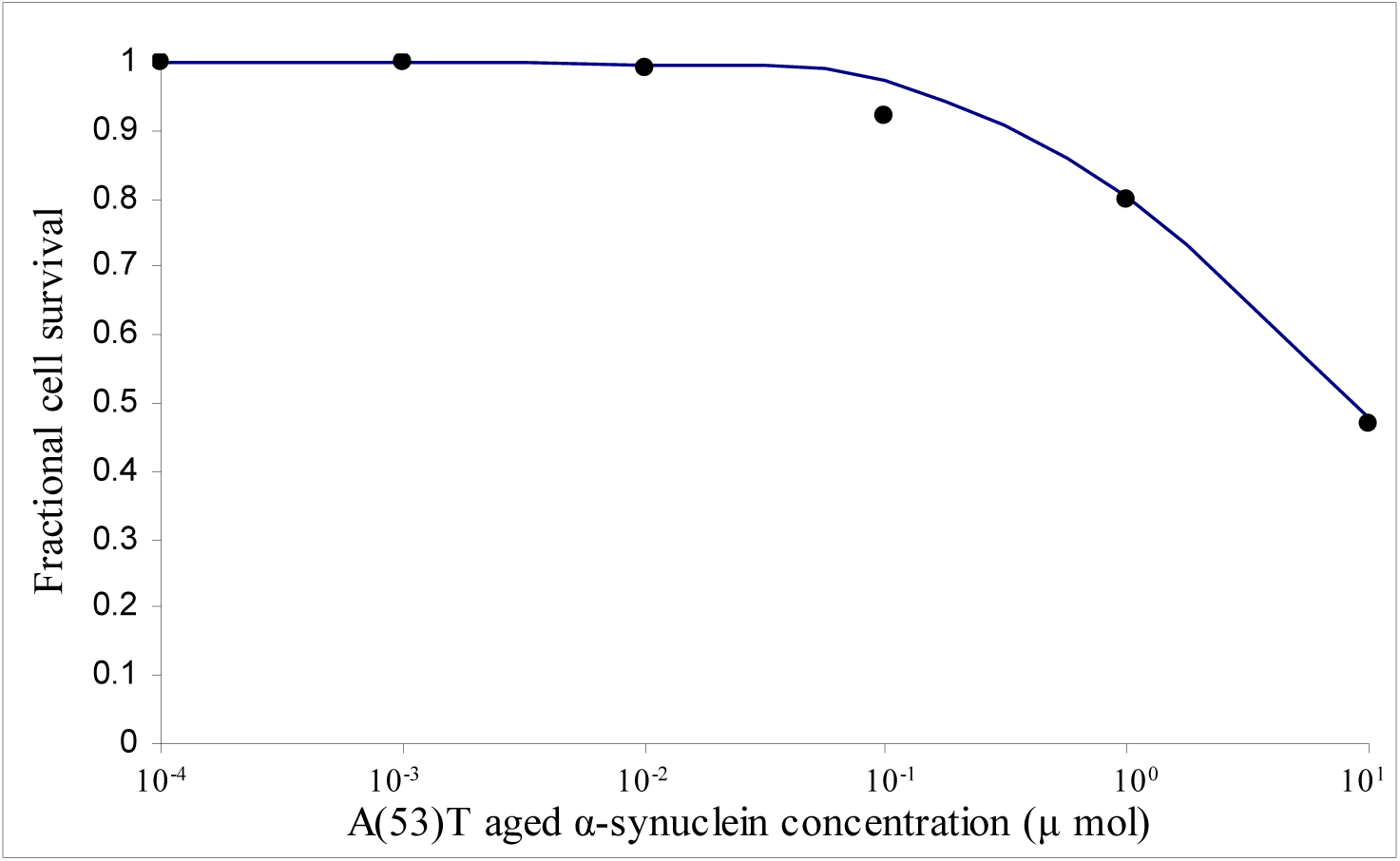
Human dopaminergic neuroblastoma SH-SY5Y cells were challenged with different concentrations of exogenous A (53)T aged α-synuclein. The experimental data (El-Agnaf et al., 1998) is compared with the results of α-synuclein Vs cell death model. (Points = Data; Continues line = Model)

**Figure 9:**
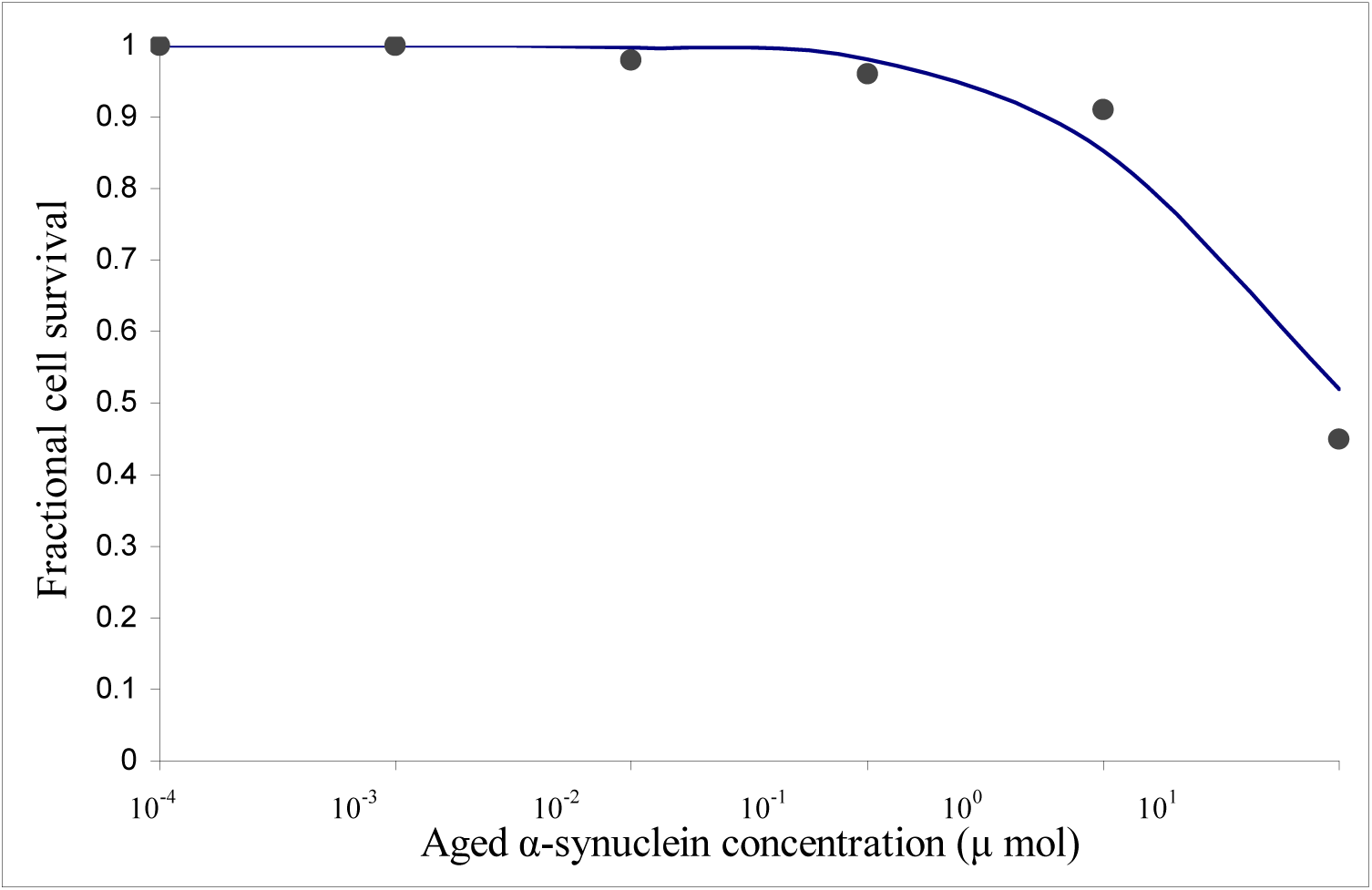
Human dopaminergic neuroblastoma SH-SY5Y cells were challenged with different concentrations of exogenous aged α-synuclein. The experimental data (El-Agnaf et al., 1998) is compared with the results of α-synuclein Vs cell death model. (Points = Data; Continues line = Model)

**Figure 10:**
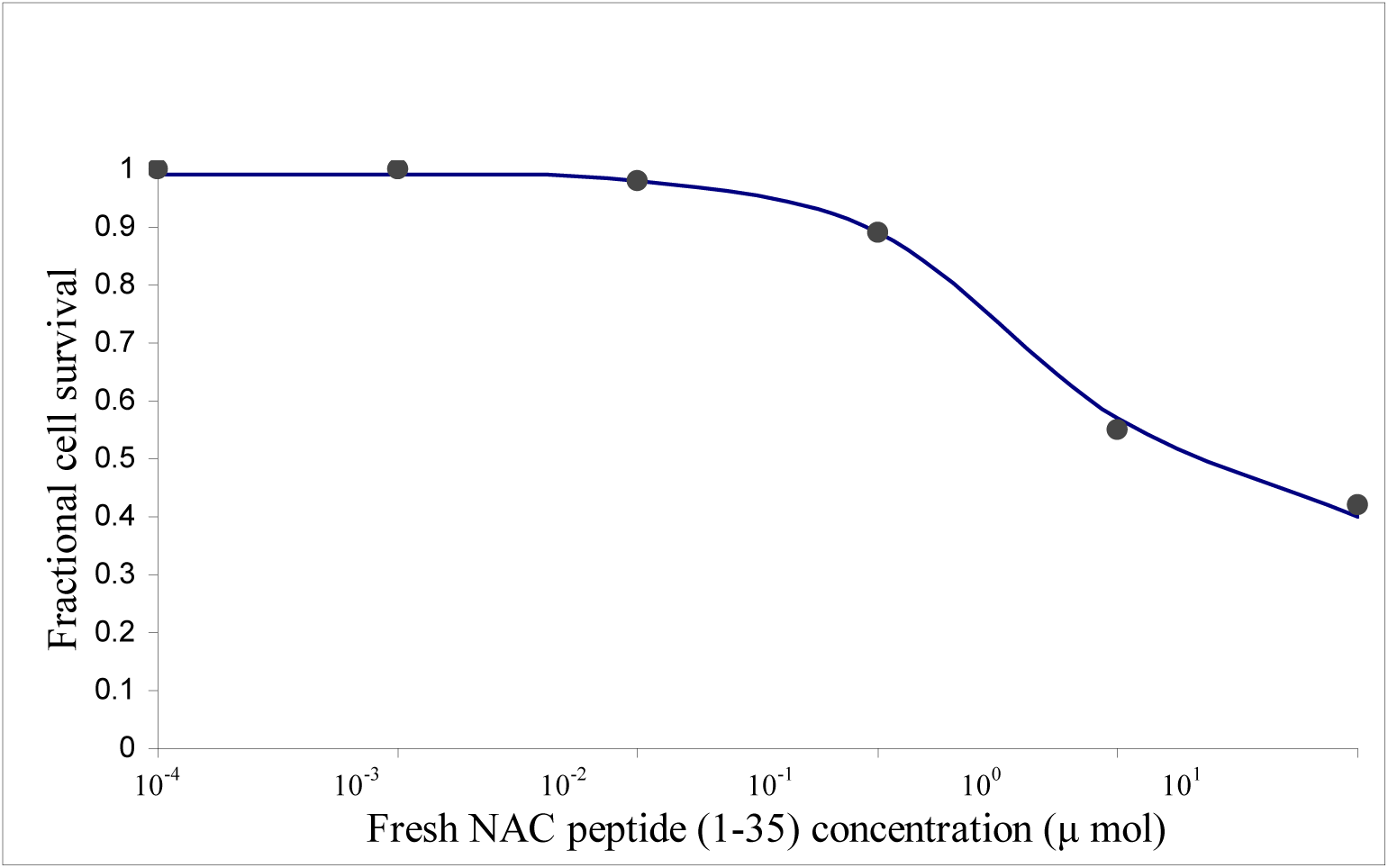
Human dopaminergic neuroblastoma SH-SY5Y cells were challenged with different concentrations of exogenous fresh NAC peptide (1-35). The experimental data (El-Agnaf et al., 1998) is compared with the results of α-synuclein Vs cell death model. (Points = Data; Continues line = Model)

**Figure 11:**
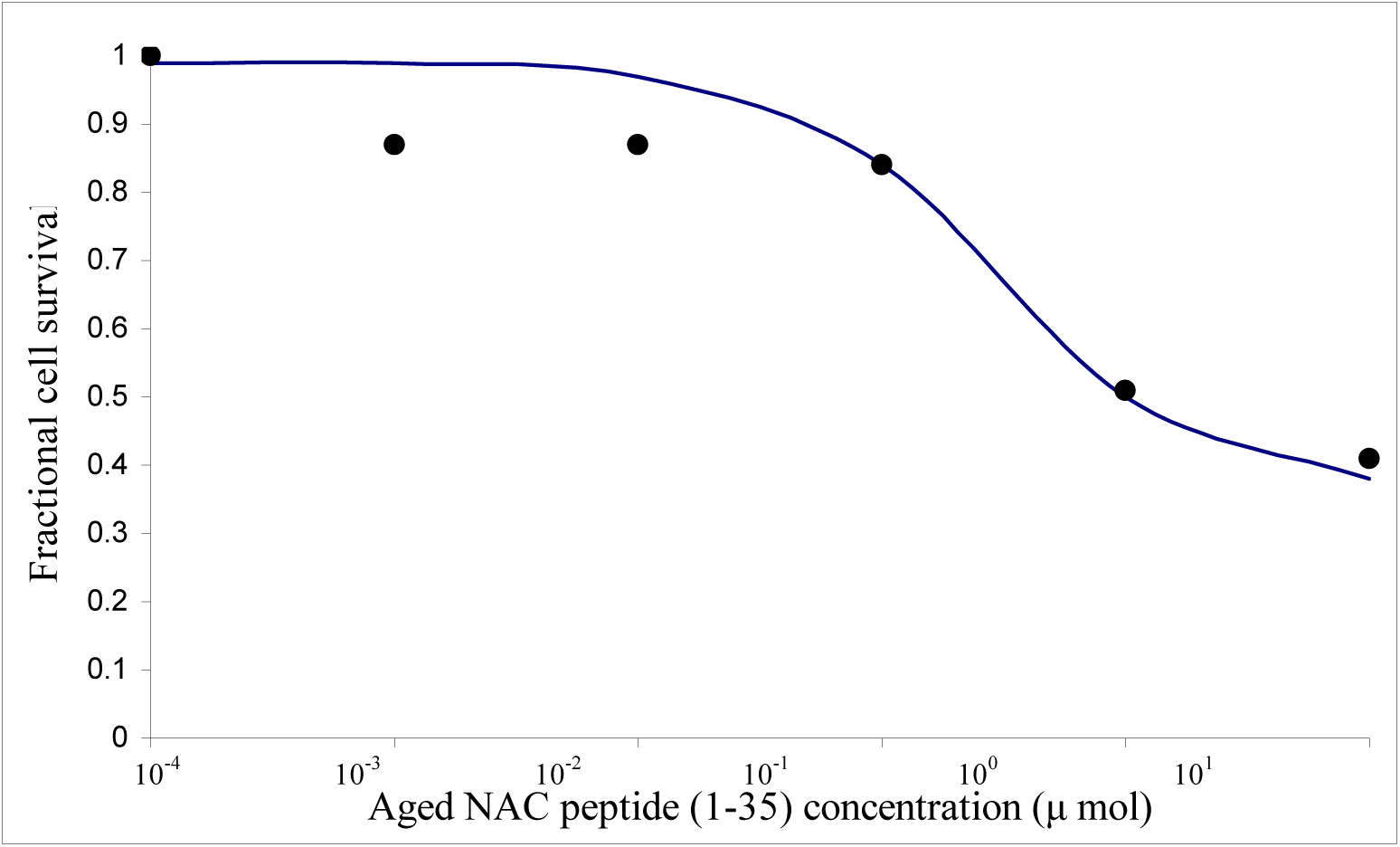
Human dopaminergic neuroblastoma SH-SY5Y cells were challenged with different concentrations of exogenous aged NAC peptide (1-35). The experimental data (El-Agnaf et al., 1998) is compared with the results of α-synuclein Vs cell death model. (Points = Data; Continues line = Model)

**Figure 12:**
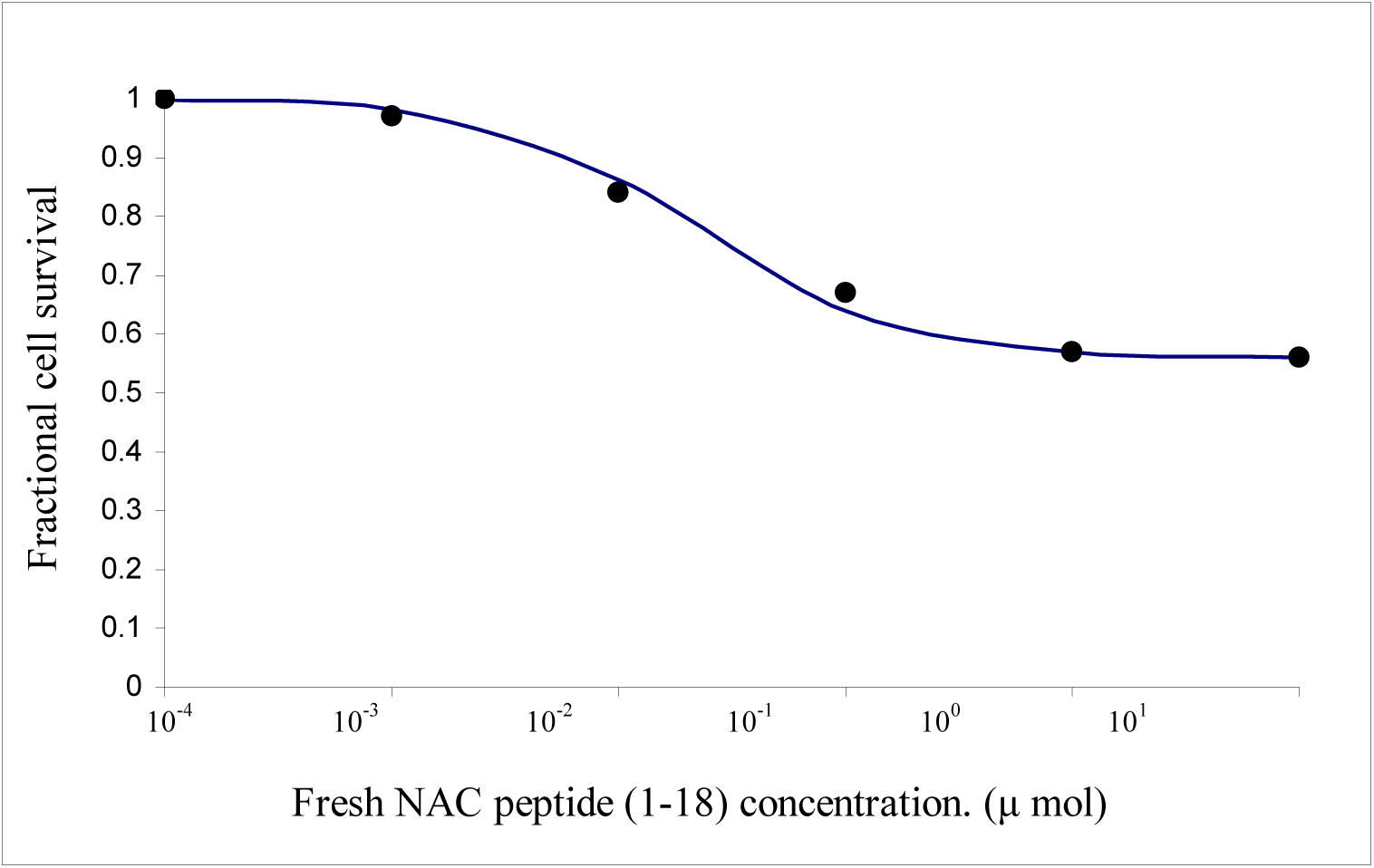
Human dopaminergic neuroblastoma SH-SY5Y cells were challenged with different concentrations of exogenous fresh NAC peptide (1-18). The experimental data (El-Agnaf et al., 1998) is compared with the results of α-synuclein Vs cell death model. (Points = Data; Continues line = Model)

**Table 1.**
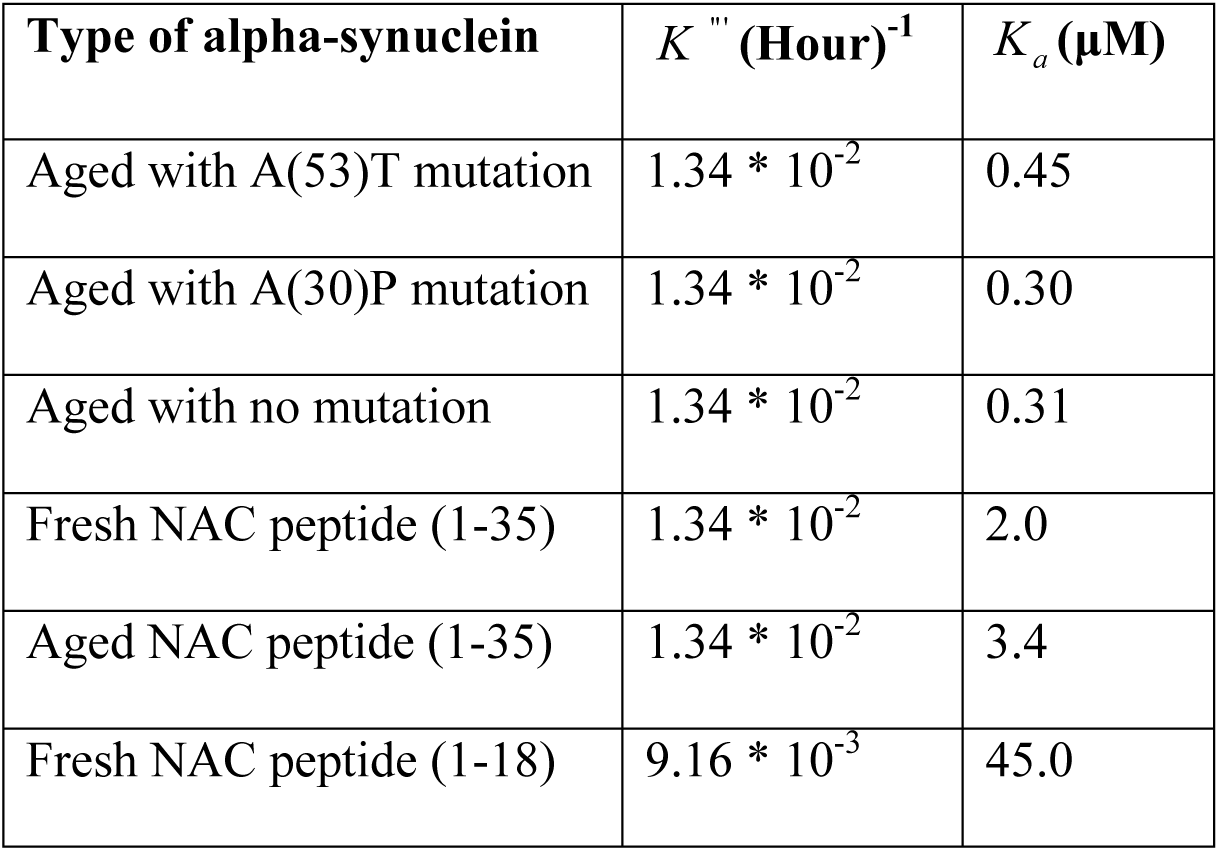
Estimated values of parameters from the experimental data.

| Type of alpha-synuclein | $K''$ (Hour)<sup>-1</sup> | $K_a$ (μM) |
| --- | --- | --- |
| Aged with A(53)T mutation | $1.34 * 10^{-2}$ | 0.45 |
| Aged with A(30)P mutation | $1.34 * 10^{-2}$ | 0.30 |
| Aged with no mutation | $1.34 * 10^{-2}$ | 0.31 |
| Fresh NAC peptide (1-35) | $1.34 * 10^{-2}$ | 2.0 |
| Aged NAC peptide (1-35) | $1.34 * 10^{-2}$ | 3.4 |
| Fresh NAC peptide (1-18) | $9.16 * 10^{-3}$ | 45.0 |

## DISCUSSION

Parkinson’s disease is a neuro degenerative disorder, with extensive loss of dopaminergic neurons. Due to this the dopamine level of the brain goes down. The remaining neurons produce more dopamine so as to compensate for the loss. This could be further pathogenic to the remaining cells. We have shown in our fractional cell survival model as a function of dopamine concentration (equation 6) that the oxidation of dopamine is autocatalytic in nature. Thus higher the intra cytoplasmic concentration of dopamine in the cell, higher is the extent of autocatalysis and higher is the accumulation of ROS. Thus, it can be concluded that once neuro degeneration starts, it increases at a much faster rate. Further it has been observed that the individuals, who respond to L-DOPA therapy in the beginning, require higher and more frequent doses later to respond. The reason for this may be that, in the beginning the body responds to L-DOPA because it is initially deficient of dopamine. L-DOPA converts into dopamine inside the brain and increases the dopamine concentration in the surrounding environment of the surviving cells. This increased concentration further increases the intra cytoplasmic concentration of dopamine. Thus it further enhances the neuro degeneration.

The reason for initiation of the disease still has not been completely established. The role of α–synuclein has always been associated with the pathogenesis of Parkinson’s disease. Two major mutations A53T & A30P have been linked with its toxicity. These substitutions have been proposed to disrupt the alpha helical structure in this region (El-Agnaf et al., 1998). This disruption may expose the highly hydrophobic NAC (61-95) region, which can initiate the aggregation of α–synuclein (Ueda et al., 1993). Recently it was shown that α–synuclein aggregation interferes with the Endoplasmic reticulum (ER)-Golgi body (GB) trafficking. It was proposed that this interference may cause the shortage of the vesicles required to sequester the dopamine being synthesized in the cytoplasm (Susan et al., 2006). The model (equation 7) has shown that the reason for cell death is not only the toxicity of α–synuclein but also the rate at which the cell produces dopamine. The *K*_*a*_ value is the measure of the toxicity of α-synuclein and the highest toxicity (*K*_*a*_) has been found from the modeling exercise for aged A53T mutated α-synuclein. This is in consistence with previous studies (Conway et al., 1998, Conway et al., 2000). The NAC peptides have been found to be much more toxic than any form of full length α–synuclein. N-terminus (1-18) region of NAC (1-35) has been identified as the amyloidogenic region, which derives the β-sheet formation and hence aggregation and deposition of NAC (1-35) (El-Agnaf et al., 1998). Consistent with this study, NAC (1-18) has been found to be highly toxic even in fresh condition (hundred times more toxic than A53T aged α-synuclein; *K*_*a*_ = 45 and 0.45 for NAC (1-18) & A(53)T aged α-synuclein respectively, see table 1). An interesting observation about this fragment is that it is not only highly toxic, but also it seems to reduce the production of dopamine and vesicles from ER. Even though the toxicity of NAC (1-18) peptide fragment is very high, the viable cell count does not come below 56% even at higher concentrations of this fragment (Figure 12).This observation could be explained by the model developed in the current study. The production of dopamine has been reduced by NAC (1-18), so the maximum dopamine that can come to the cytoplasm when no vesicle is present to sequester it (the condition generated at high concentration of α-synuclein), is less. Thus the generation of ROS is low and so the cell degeneration is also less. But still the symptoms of Parkinson’s disease will be generated in the animal because the ultimate reason of Parkinsonism is the deficiency of dopamine, which is reduced directly by NAC (1-18) peptide.

*K*_*a*_ & *K*‴ are the two constants in the model (equation 7) which determine the shape of the curve. *K*_*a*_ gives the information about the toxicity of various types of α– synuclein, whereas *K*‴ tells about the rate of production of vesicles from Endoplasmic Reticulum and also the production of dopamine in the cytoplasm (Figure.1). *K*_*a*_ represents the tendency of α–synuclein to inhibit the ER-GB trafficking. Higher the value of *K*_*a*_, lower is the concentration of α–synuclein at which it will hinder almost all the vesicles form ER to fuse with GB. Thus at lower concentration of α–synuclein almost all the dopamine formed in the cytoplasm will be left free. This will cause the viable cell count to fall almost immediately. Thus the higher *K*_*a*_ value shifts the curve towards left and vice versa (See figure 13).

**Figure 13:**
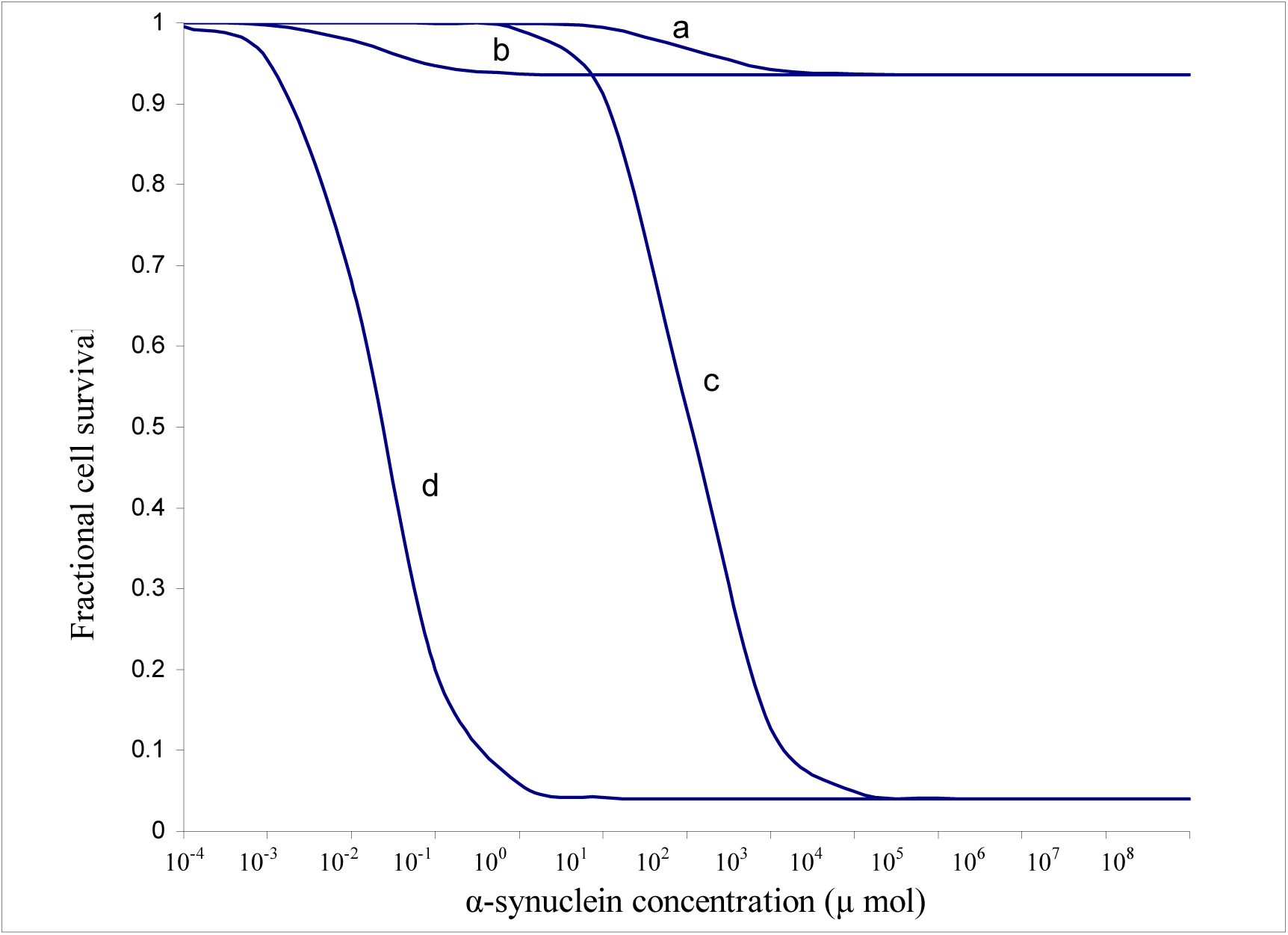
Simulation of α-synuclein Vs cell death model with different values of *K* and *K*_*a*_.

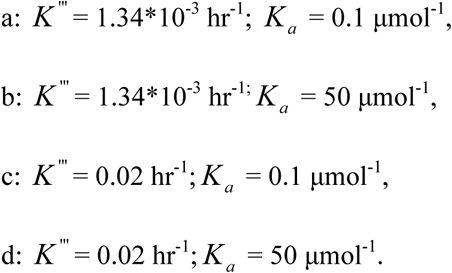

*K*‴ represents the capacity of the cell to produce dopamine. Higher the value of *K*‴ higher is the rate of production of dopamine, thus higher is the concentration of dopamine in the cytoplasm. In normal healthy cells the rate of production of dopamine in dopaminergic cells is matched with the higher rate of production of vesicles from ER, hence there is no accumulation of dopamine in the cytoplasm. In diseased cells the abnormal α–synuclein hinders the ER-GB trafficking and thus decreases the vesicles required to sequester the dopamine from the cytoplasm. Thus at very high concentrations of α–synuclein, the cytoplasmic concentration of dopamine will become very high, reducing the viable cell count to very low value. Higher *K*‴ value shifts the curve down and vice versa.

A mathematical model has been developed relating pathogenic α–synuclein and dopamine amount to cell death. Based on this analysis we have been able to conclude that oxidation of dopamine to produce ROS is an autocatalytic step. A53T mutation is more toxic than either A30P or WT α–synuclein, which is consistent with the experimental studies. (Conway et al., 1998; Conway et al., 2000). Further NAC peptide is highly pathogenic and, not only it hinders the early secretary pathway, but also it reduces the dopamine production in the cells.

The conversion of normal α–synuclein to fibrilised form due to the action of ROS has not been studied in this paper since no data has been reported in literature relating these two species. The conversion of healthy cells to dead cells may go through the formation of lewy bodies, DNA damage, peroxidation of proteins etc, but the model does not include these aspects.

## ABBREVIATIONS

AD: Alzheimer’s disease.
ER: Endoplasmic Reticulum
GB: Golgi body
H_2_O_2_: Hydrogen peroxide.
NAC: Non-Aβ Component of Alzheimer’s disease.
6-OHDA: 6-hydroxy dopamine.
PD: Parkinson’s disease
ROS: Reactive Oxygen Species
VMAT2: Vesicular mono amine transporter 2

## APPENDIX

### Assumptions

1. In a living cell all the metabolites remain at steady state concentration (except those metabolites which are affected by signal transduction process). Thus, we have taken four steady state points in our model.
2. The two reactions i.e. generation of vesicles from endoplasmic reticulum[ER] and synthesis of dopamine have been assumed to be of zero order reactions
3. The α-synuclein is taken in excess, thus concentration of α-synuclein does not reduce considerably with respect to time.
4. The cell death due to ROS has been observed to be dependent on ROS concentration alone and not on the initial cell concentration of healthy cells (Kanda et al. (2000)). Thus this reaction is taken as pseudo first order reaction

### A. Derivation for Dopamine Vs cell death model

Let [D] = Dopamine; [ROS] =Reactive Oxygen Species;

[C] = Initial cell concentration; [DC] = Dead cell concentration.

We wrote the mechanism as:

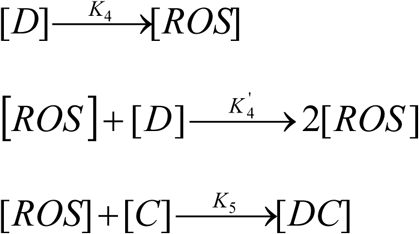

Steady state approximation:

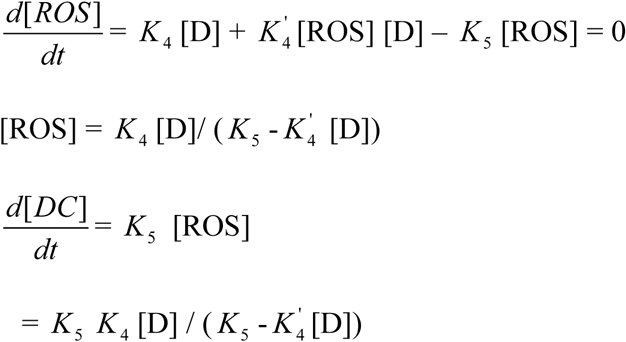

Now since 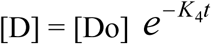

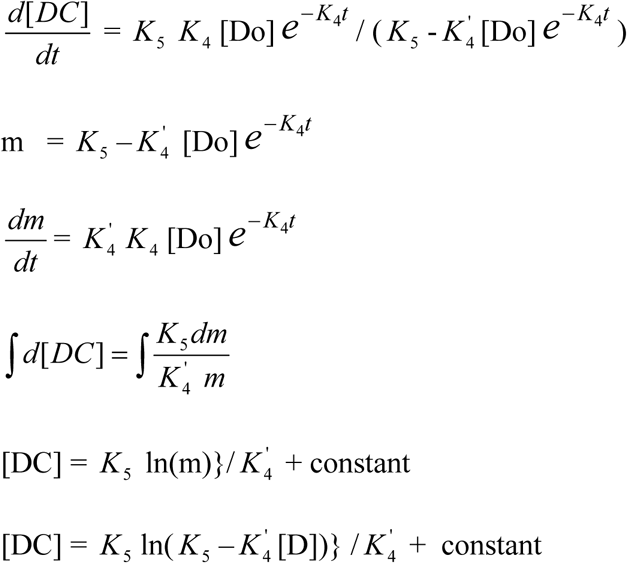

At t = 0 [DC] = 0 & [D] = [Do]

So 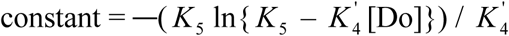

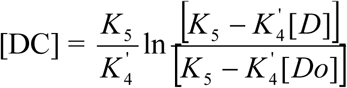

But at steady state [D] = 0

So 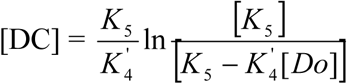

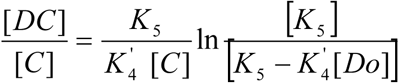

Subtracting both sides from one we get:

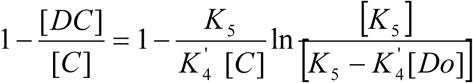

Let 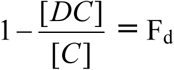 (Fractional cell survival), 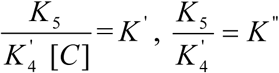

Thus 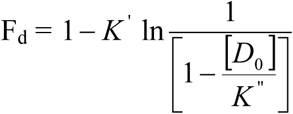

Where *K*′ is unit less constant and [C] is the concentration of cells in the beginning.

### B. Derivation for effect of antioxidant on cell death

From steady state approximation we get:

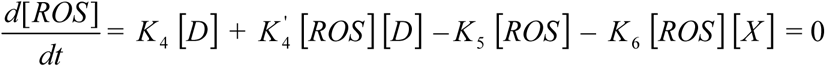

Where [*X*] = concentration of anti-oxidant

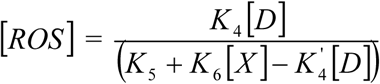

This on proceeding as before gives:

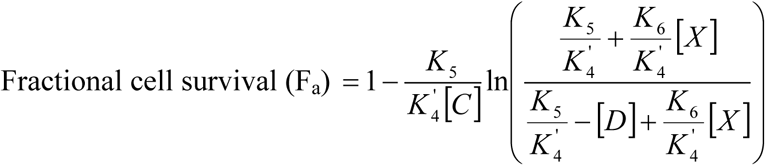

Where [*C*] is the initial cell concentration.

Let 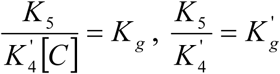 and 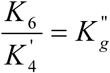

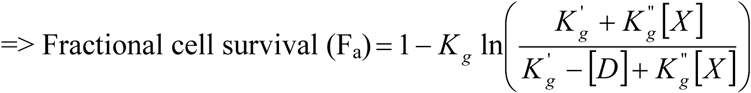

### C. Derivation for alpha-synuclein Vs cell death model (Figure 3)

Let:

[V] = Concentration of Vesicles emerging from Endoplasmic Reticulum(ER).

[A] = Concentration of Pathological alpha-Synuclein.

[V_s_] = Concentration of Vesicles emerging from Golgi body (GB), committed to accept free Dopamine, being synthesized in cytoplasm.

[D] = Concentration of Dopamine synthesizing in the cytoplasm.

[ROS] = Reactive Oxygen Species (H_2_O_2_) produced due to the oxidation of dopamine.

[DC] = Dead Cell Concentration.

[C] = Initial cell concentration.

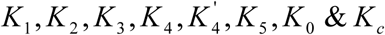 are Rate constants of different reactions.

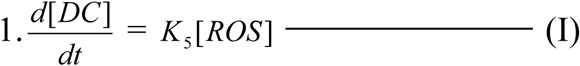

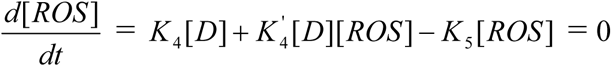 (Steady state approximations.)

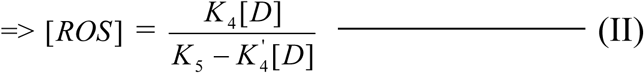

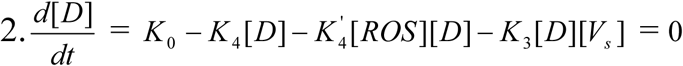

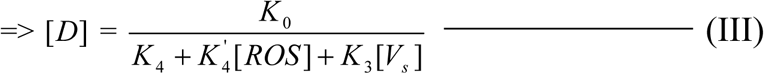

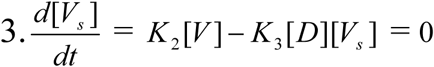

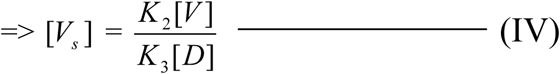

Putting values of [*ROS*] & [*V*_*s*_] in equation (III) we get:

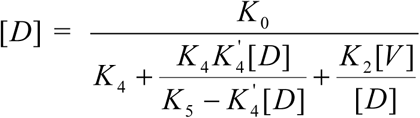

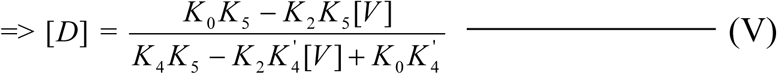

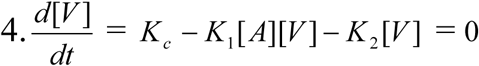

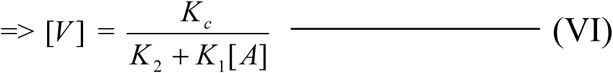

Putting the value of [*V*] in equation (V) we get:

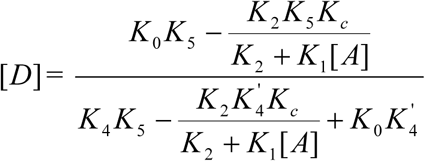

Putting the value of [*D*] in equation (II) and solving for[*ROS*] we get:

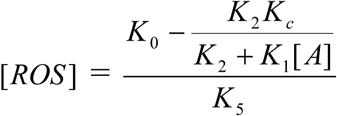

Putting the value of [*ROS*] in equation (I) and integrating for[*DC*] we get:

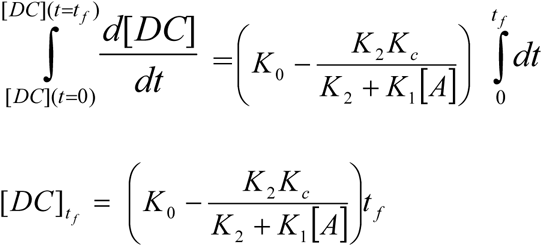

Now when 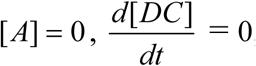

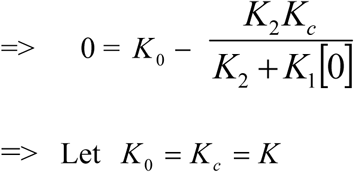

Thus

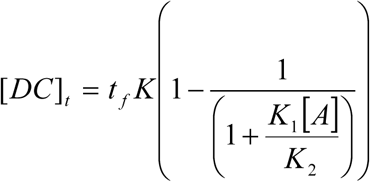

Dividing both sides by initial cell concentration [*C*] & subtracting both sides from one we get:

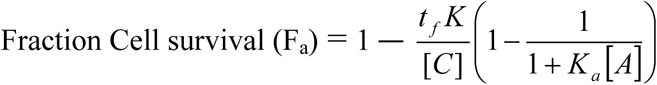

Let 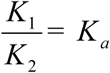 and 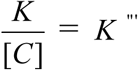

Thus 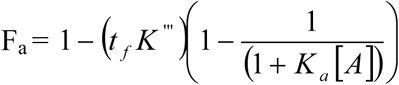

Where *K*‴ and *K*_*a*_ are constants with units time^−1^ and (µ mol)^−1^ respectively.

